# Using SuperClomeleon to measure changes in intracellular chloride during development and after early life stress

**DOI:** 10.1101/2022.10.06.511115

**Authors:** L.J. Herstel, C. Peerboom, S. Uijtewaal, D. Selemangel, H. Karst, C.J. Wierenga

## Abstract

Intraneuronal chloride concentrations ([Cl^-^]_i_) decrease during development resulting in a shift from depolarizing to hyperpolarizing γ-aminobutyric acid (GABA) responses via chloride-permeable GABA_A_ receptors. This GABA shift plays a pivotal role in postnatal brain development, and can be strongly influenced by early life experience. Here, we assessed the applicability of the recently developed fluorescent SuperClomeleon (SClm) sensor to examine changes in [Cl^-^]_i_ using two-photon microscopy in brain slices. We used SClm mice of both sexes to monitor the developmental decrease in neuronal chloride levels in organotypic hippocampal cultures. We could discern a clear reduction in [Cl^-^]_i_ between DIV3 and DIV9 (equivalent to the second postnatal week *in vivo)* and a further decrease in some cells until DIV22. In addition, we assessed alterations in [Cl^-^]_i_ in the medial prefrontal cortex (mPFC) of P9 male SClm mouse pups after early life stress (ELS). ELS was induced by limiting nesting material between P2 and P9. ELS induced a shift towards higher (i.e. immature) chloride levels in layer 2/3 cells in the mPFC. Although conversion from SClm fluorescence to absolute chloride concentrations proved difficult, our study underscores that the SClm sensor is a powerful tool to measure physiological changes in [Cl^-^]_i_ in brain slices.

**Significance Statement:** The reduction of intraneuronal chloride concentrations is crucial for brain development, as it ensures a shift from the initial excitatory action of the neurotransmitter GABA in immature neurons to the inhibitory GABA signaling in the adult brain. Despite the significance of chloride maturation, it has been difficult to study this phenomenon in experiments. Recent development of chloride sensors enable direct imaging of intracellular chloride signaling in neurons. Here we assessed the applicability of the SuperClomeleon chloride sensor to measure physiologically relevant changes in chloride levels using two-photon microscopy in cultured and acute brain slices. Although we also point out some limitations, we conclude that the SuperClomeleon sensor is a powerful tool to measure physiological changes in intracellular chloride.

## Introduction

During normal neuronal development, γ-aminobutyric acid (GABA) responses through ionotropic GABA_A_ receptors shift from depolarizing to hyperpolarizing as a result of the developmental decrease in intracellular chloride concentration. In the immature brain, the intracellular chloride concentration is high and activation of GABA_A_ receptors results in an outflow of chloride leading to membrane depolarization. During early postnatal development, intracellular chloride levels gradually decrease. As a result, activation of GABA_A_ receptors in mature neurons leads to the influx of chloride and GABAergic signaling induces membrane hyperpolarization (Ben-Ari et al., 2007; Kaila et al., 2014; Rivera et al., 1999). This shift in GABA signaling plays a pivotal role in postnatal neuronal development and its timing affects brain function throughout life (Kaila et al., 2014; Lohmann and Kessels, 2014; Peerboom and Wierenga, 2021; Sernagor et al., 2010).

In rodents the GABA shift occurs normally between postnatal day (P)10 and 14 depending on brain region and cell type (Ben-Ari et al., 2007; Glykys et al., 2009; Kirmse et al., 2015; Rivera et al., 1999; Romo-Parra et al., 2008; Stein et al., 2004; Sulis Sato et al., 2017). For example, intracellular chloride levels in the visual cortex mature several days earlier compared to the hippocampus (Murata and Colonnese, 2020), while in the prefrontal cortex the GABA shift occurs even later (Amadeo et al., 2018; Karst et al., 2019). In addition, GABAergic maturation has been shown to be strongly influenced by experiences during early life. For instance, prenatal maternal restraint stress as well as repeated separations of newborn pups from their mother induced a delay in the GABA shift in hippocampal pyramidal cells in young mice (Furukawa et al., 2017; Hu et al., 2017; Veerawatananan et al., 2016). Early life stress (ELS) has life-long consequences on neurophysiology and behavior in both humans and rodents and poses an increased risk for psychopathology later in life (Joëls et al., 2018; Teicher et al., 2016). The medial prefrontal cortex (mPFC) is known to be very sensitive to stress early in life, with life-long consequences for anxiety and stress responses (Ishikawa et al., 2015; Karst et al., 2020). The mPFC functions as a central coordinator of stress responses across brain regions as well as the periphery (McKlveen et al., 2015). However, it is currently unclear how GABA signaling in the mPFC is affected by ELS.

In most studies intracellular chloride concentrations in neurons are determined using perforated patch clamp recordings. With this technique antibiotics (e.g. gramicidin or amphotericin B) are included in the pipette to form small pores in the membrane which leaves intracellular chloride concentration intact. However, perforated patch clamp recordings are time intensive and it is difficult to perform long recordings, as access is not stable (Arosio et al., 2010). In addition, large individual differences can exist between neurons and many individual recordings may be required to get a good population estimate (Sulis Sato et al., 2017; Tyzio et al., 2007). As a promising alternative, biosensors are being developed which allow for the real time measurement of intracellular chloride levels in a noninvasive manner (Arosio et al., 2010). The SuperClomeleon (SClm) sensor (Grimley et al., 2013) is a second generation chloride sensor with chloride sensitivity in the physiological range. The SClm sensor consists of two fluorescent proteins, Cerulean (CFP mutant) and Topaz (YFP mutant), joined by a flexible linker. Depending on the binding of chloride, Fluorescence Resonance Energy Transfer (FRET) occurs from the CFP donor to the YFP acceptor (Grimley et al., 2013). FRET ratios (fluorescence intensity of YFP/CFP) are independent of expression level and imaging settings, which is a major advantage when imaging in intact brain tissue. The SClm sensor has successfully been used to determine the steady-state intracellular chloride concentration in adult mice *in vivo* (Boffi et al., 2018) and to demonstrate the existence of cytoplasmic chloride microdomains in neurons (Rahmati et al., 2021), but it has not been used to examine chloride maturation during neuronal development.

Here, we used the SClm sensor to detect changes in chloride during early postnatal development in organotypic hippocampal cultures of mice and in acute slices of the prefrontal cortex from young control mice and mice who experienced ELS.

## Methods

### Animals

SuperClomeleon^lox/-^ mice (Rahmati et al., 2021) (a gift from Kevin Staley, Massachusetts General Hospital, Boston, MA) were crossed with CamKIIα^Cre/-^ mice (Casanova et al., 2001; Tsien et al., 1996) (a gift from Stefan Berger, German Cancer Research Center, Heidelberg, Germany) and will hereafter be referred to as SClm mice. Animals were housed at reversed day-night cycle with a room temperature of 22 ± 2 °C and humidity of approximately 65%. Food (standard chow) and water were provided *ad libitum.* We noticed that SClm mice had poor breeding performance compared to other strains kept in the same facility. On postnatal day (P) 2 the litter was randomly assigned to either the control condition (standard housing) or the ELS condition. In the ELS condition a limited amount of nesting and bedding material was made available between P2 and P9 (Karst et al., 2020; Naninck et al., 2015; Rice et al., 2008). All animal experiments were performed in compliance with the guidelines for the welfare of experimental animals and were approved by the local authorities.

### Organotypic culture preparation

Postnatal developmental changes in intracellular chloride concentration ([Cl^-^]_i_) were imaged in organotypic hippocampal cultures made from P6 control SClm mice of both sexes. For the chloride calibration and wash-in experiments, organotypic hippocampal cultures were made from P6 WT C57BL/6 mice and SClm expression was achieved by viral injection. Slice cultures were prepared using a method based on Stoppini *et al.* (1991). After decapitation the brain was rapidly removed and placed in ice-cold Gey’s Balanced Salt Solution (GBSS; containing (in mM): 137 NaCl, 5 KCl, 1.5 CaCl_2_, 1 MgCl_2_, 0.3 MgSO_4_, 0.2 KH_2_PO_4_ and 0.85 Na_2_HPO_4_) with 25 mM glucose, 12.5 mM HEPES and 1 mM kynurenic acid. Transverse hippocampal slices of 400 μm thick were cut with a McIlwain tissue chopper (Brinkmann Instruments). Slices were placed on Millicell membrane inserts (Millipore) in wells containing 1 mL culture medium (consisting of 48% MEM, 25% HBSS, 25% horse serum, 25 mM glucose, and 12.5 mM HEPES, with an osmolarity of 325 mOsm and a pH of 7.3 – 7.4). Slices were stored in an incubator (35°C with 5% CO_2_) and medium was replayed three times a week. Experiments were performed after 1 to 22 days in vitro (DIV).

### Viral expression

An adeno-associated virus (AAV) with Cre-independent expression of SClm under the control of the Synapsin promotor (AAV9.hSyn.sCLM; a gift from Kevin Staley, Massachusetts General Hospital, Boston, MA) was injected in the CA1 area of WT cultured hippocampal slices on DIV1. Slices were imaged on DIV9-16. Compared to slices from SClm mice, viral expression of the SClm sensor resulted in larger variability in neuronal FRET (YFP/CFP) ratios, probably due to variability in slice quality and viral expression levels. We optimized the viral concentration to get comparable levels of YFP and CFP fluorescent intensity as observed in the mouse line.

### Acute slice preparation

Young SClm mice were decapitated at P9, followed by quick removal of the brain. For this study we used only male mice to allow direct comparison with our earlier study in C57/BL6 mice (Karst et al., 2019). Mice were always decapitated in the morning to eliminate influences of fluctuating corticosterone (CORT) levels during the day, due to the reversed day-night cycle this means that CORT levels are high at that time. The brain was stored in ice cold artificial cerebrospinal fluid (ACSF, containing (in mM): 120 choline chloride, 3.5 KCl, 0.5 CaCl_2_, 6 MgSO_4_, 1.25 NaH_2_PO_4_, 25 D-glucose and 25 NaHCO_3_). Coronal slices of 350 μm thickness were made with a vibratome (Leica VT 1000S). After placing them in ACSF (consisting of (in mM): 120 NaCl, 3.5 KCl, 1.3 MgSO_4_, 1.25 NaH_2_PO_4_, 2.5 CaCl_2_, 25 D-glucose and 25 NaHCO_3_) slices were heat shocked at 32°C for 20 min. Slices were then kept at room temperature and after recovery for at least 1 h, transported individually in Eppendorf tubes filled with ACSF to the microscope room in another building with a transportation time of 10 minutes.

### Two-photon imaging

Slices were transferred to the microscope chamber. The bath was continuously perfused with carbonated (95% O2, 5% CO_2_) ACSF (in mM: 126 NaCl, 3 KCl, 2.5 CaCl_2_, 1.3 MgCl_2_, 26 NaHCO_3_, 1.25 NaH_2_PO_4_, 20 D-glucose and 1 Trolox, with an osmolarity of 310 ± 10 mOsm/L) at a rate of approximately 1 mL/min. Bath temperature was monitored and maintained at 30–32 °C throughout the experiment. Two-photon imaging of pyramidal neurons in layer 2/3 of the mPFC or pyramidal neurons in the CA1 area of the hippocampus was performed on a customized two-photon laser scanning microscope (Femto2D, Femtonics, Budapest, Hungary). To excite the CFP donor, a Ti-Sapphire femtosecond pulsed laser (MaiTai HP, Spectra-Physics) was tuned to 840 nm. The emission signal was split using a dichroic beam splitter at 505 nm and detected using two GaAsP photomultiplier tubes. We collected fluorescence emission of Cerulean/CFP (485 ± 15 nm) and Topaz/YFP (535 ± 15 nm) in parallel. A 60x water immersion objective (Nikon NIR Apochromat; NA 1.0) was used to locate the cell layer. Of each slice, 2-4 image z-stacks in different field of views (FOVs) were acquired at a resolution of 8.1 pixels/μm (1024×1024 pixels, 126×126 μm) with 1 μm steps of approximately 40-85 μm in depth. To monitor acute changes in [Cl^-^]_i_ we bath applied GABA_A_ receptor agonist muscimol (Tocris, 10 μM) and imaged at lower resolution (every 2 minutes at a resolution of 4.1 pixels/μm (512×512 pixels, 126×126 μm) with 1 μm steps of 30-50 μm in depth).

### Imaging data analysis

Image analysis was performed using Fiji/ImageJ software and results were analyzed in Prism 9 (GraphPad). We manually determined regions of interest (ROIs) around individual neuron somata. To analyze a representative cell population, in each image z-stack we selected four z-planes at comparable depths in which three cells were identified that varied in brightness (bright, middle and dark). We subtracted the mean fluorescence intensity of the background in the same image plane from the mean fluorescence intensity of CFP and YFP before calculating the fluorescence ratio. We limited our analysis to cells which were located within 450 pixels from the center of the image, as FRET ratios showed slight aberrations at the edge of our images. We excluded cells with a FRET ratio < 0.5 or > 1.6, to avoid the selection of unhealthy cells (59 of 1893 cells; 3.1%). We verified that inclusion of these cells did not change our conclusions. We confirmed that the FRET ratios of individual cells were uncorrelated with their fluorescence intensities.

FRET-colored images (as shown in Fig. 1C, 2D, 3A and 4A) were made in ImageJ. We first subtracted the average background and Gaussian filtered the CFP and YFP image z-stacks separately. Next, the acceptor (YFP) image was divided by the donor (CFP) image to get the ratiometric image. A mask was created by manually drawing ROIs for each soma in the image. An average projection of the radiometric image was made of a specific z range and multiplied by the mask. Finally, the masked ratiometric image was combined with the grayscale image.

**Figure 1.**
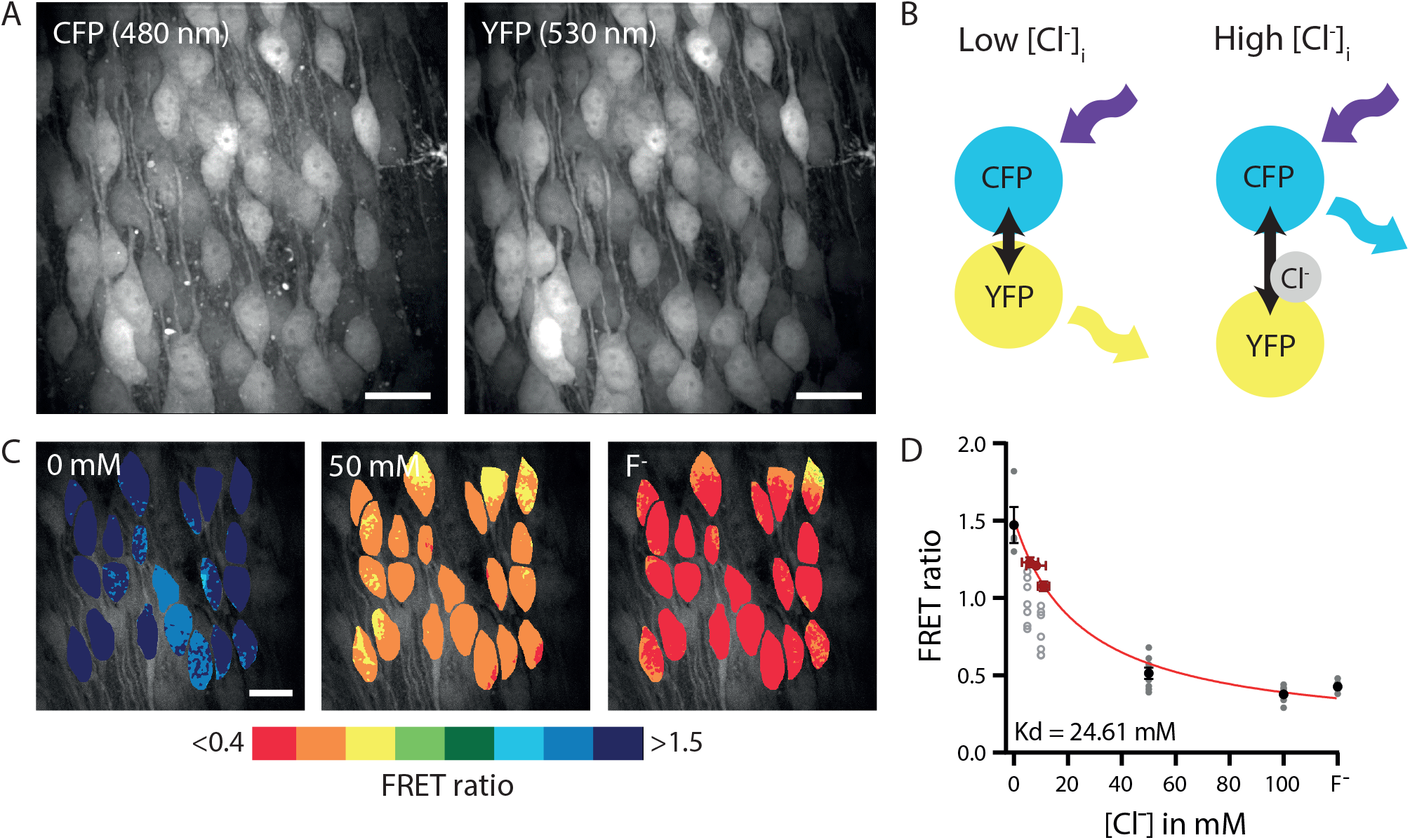
Two-photon imaging of [Cl^-^]_i_ in brain slices. A. Example image of CFP (480 nm) and YFP (530 nm) fluorescence in an organotypic hippocampal culture from a SClm mouse. Scale bar: 20 μm. B. Illustration of Fluorescence Resonance Energy Transfer (FRET) from CFP donor to YFP acceptor of the SClm sensor. FRET values (YFP/CFP fluorescence ratio) decrease with higher chloride concentrations. C. Two-photon imaging of chloride-dependent changes in the FRET ratio (530/480 nm emission) in a WT organotypic hippocampal culture with AAV SClm expression. [Cl^-^]_i_ was clamped to the indicated external chloride concentration via ionophore treatment. Individual cells are color-coded to their FRET ratios. Scale bar: 20 μm. D. Calibration curve constructed from ionophore experiments (black/grey symbols) and perforated patch (red symbols) data. Data is presented as mean ± SEM. These data was fit by equation (2), yielding the following fit parameters: K_d_ = 24.6 mM, R_max_ = 1.51, R_min_ = 0.12 (see methods for details). This calibration curve was used to convert FRET ratios into estimated chloride levels in the rest of this study. We also show the individual data points representing individual ionophore experiments (average over 12 cells per experiment). At 5 and 10 mM extracellular chloride FRET values were highly variable (open symbols). As described in the methods, we excluded these data points from our analysis and resorted to perforated patch recordings (red symbols) for this chloride range. The grey dotted line shows the alternative calibration curve when all ionophore data are included (fit parameters K_d_ = 8.4; R_max_ = 1.35; R_min_ = 0.32; without perforated patch data).

**Figure 2.**
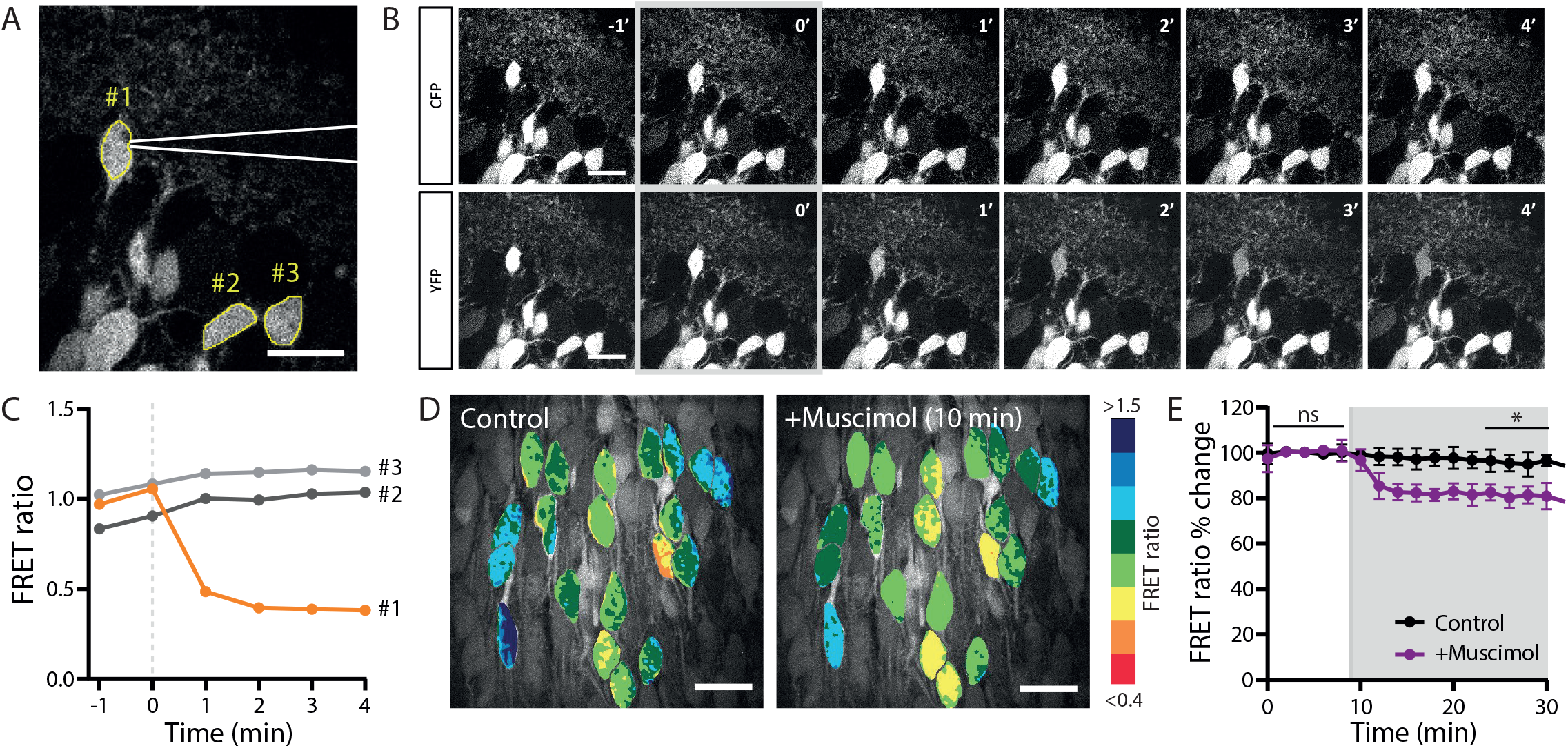
Monitoring acute changes in [Cl^-^]_i_ with SClm. A. Two-photon image of CA1 pyramidal neurons in the hippocampus of an acute slice from a SClm mouse. A patch pipette (in white) is attached to cell #1 for a whole-cell recording. Two control cells are indicated with #2 and #3. Scale bar: 20 μm. B. Time course of CFP (upper row) and YFP (lower row) fluorescence right before and during the first minutes after break-in. After break-in (0’; gray) cell #1 rapidly fills with the high chloride internal solution (70 mM KCl) resulting in a decrease in YFP fluorescence. Scale bar: 20 μm. C. Corresponding FRET ratio for the two-photon images in B for the patched cell (#1) and control cells (#2 and #3) over time. D. Acute wash-in with muscimol in cultured slices with viral SClm expression, caused a decrease in FRET ratio in CA1 pyramidal neurons within 10 minutes. Scale bar: 20 μm. E. Average FRET ratios over time during wash-in of muscimol (grey area) and control. Data from 84 cells, 7 slices, 4 mice in both groups.

**Figure 3.**
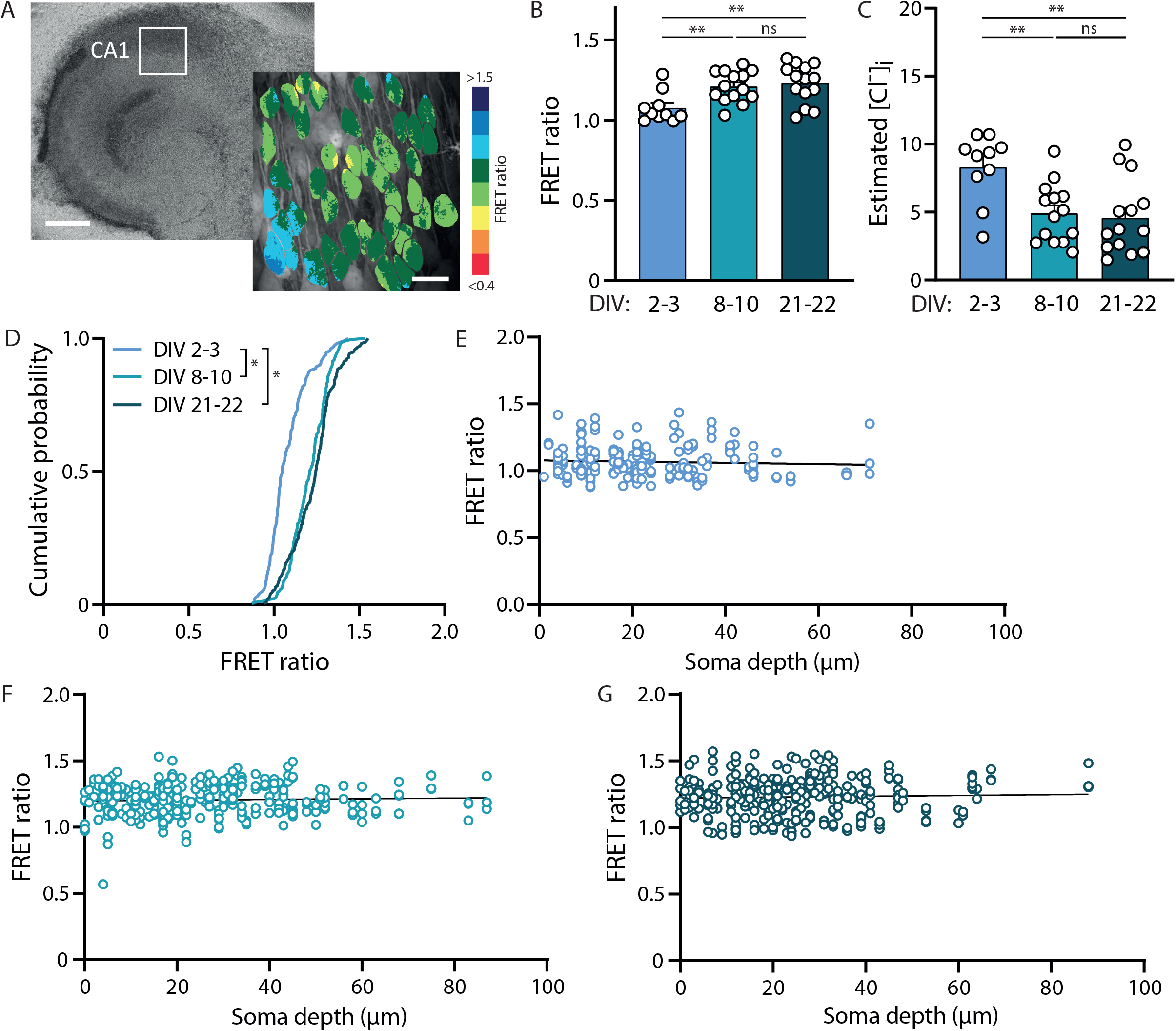
Developmental decrease in [Cl^-^]_i_ continues in organotypic cultures. A. Example of an organotypic hippocampal culture from a SClm mouse at DIV8. Scale bar: 500 μm. In the zoom an example of the FRET ratios determined in the CA1 area. Scale bar: 20 μm. B. Average FRET ratios of CA1 pyramidal cells at DIV2-3 (n = 10 slices of 6 mice), DIV8-10 (n = 14 slices of 8 mice) and DIV21-22 (n = 14 slices of 8 mice). There was a significant increase in FRET ratio over time (p = 0.033 DIV2-3 vs DIV8-10; p = 0.008 DIV2-3 vs DIV21-22; p > 0.99 DIV8-10 vs DIV21-22; KW test). C. Average estimated [Cl^-^]_i_ as calculated from the FRET ratios in B. There was a significant decrease in [Cl^-^]_i_ over time (p = 0.006 DIV2-3 vs DIV8-10; p =0.003 DIV2-3 vs DIV21-22; p = 0.93 DIV8-10 vs DIV21-22; one-way ANOVA). D. Cumulative distribution of FRET ratios for individual cells at the 3 time points. For each slice 15 cells were randomly selected, making the total of number of plotted cells per condition at least 150 (p < 0.001 DIV2-3 vs. DIV8-10 and vs. DIV21-22; p = 0.053 DIV8-10 vs. DIV21-22; KS test). E. Individual FRET ratios plotted against soma depth in slices at DIV2-3 (n = 186 neurons). Line represents linear regression fit (r = 0.004; p = 0.41). F. Same as E at DIV8-10 (n = 299 neurons). Line represents linear regression fit (r = 0.002; p = 0.48). G. Same as E at DIV21-22 (n = 314 neurons). Line represents linear regression fit (r = 0.001; p = 0.50).

**Figure 4.**
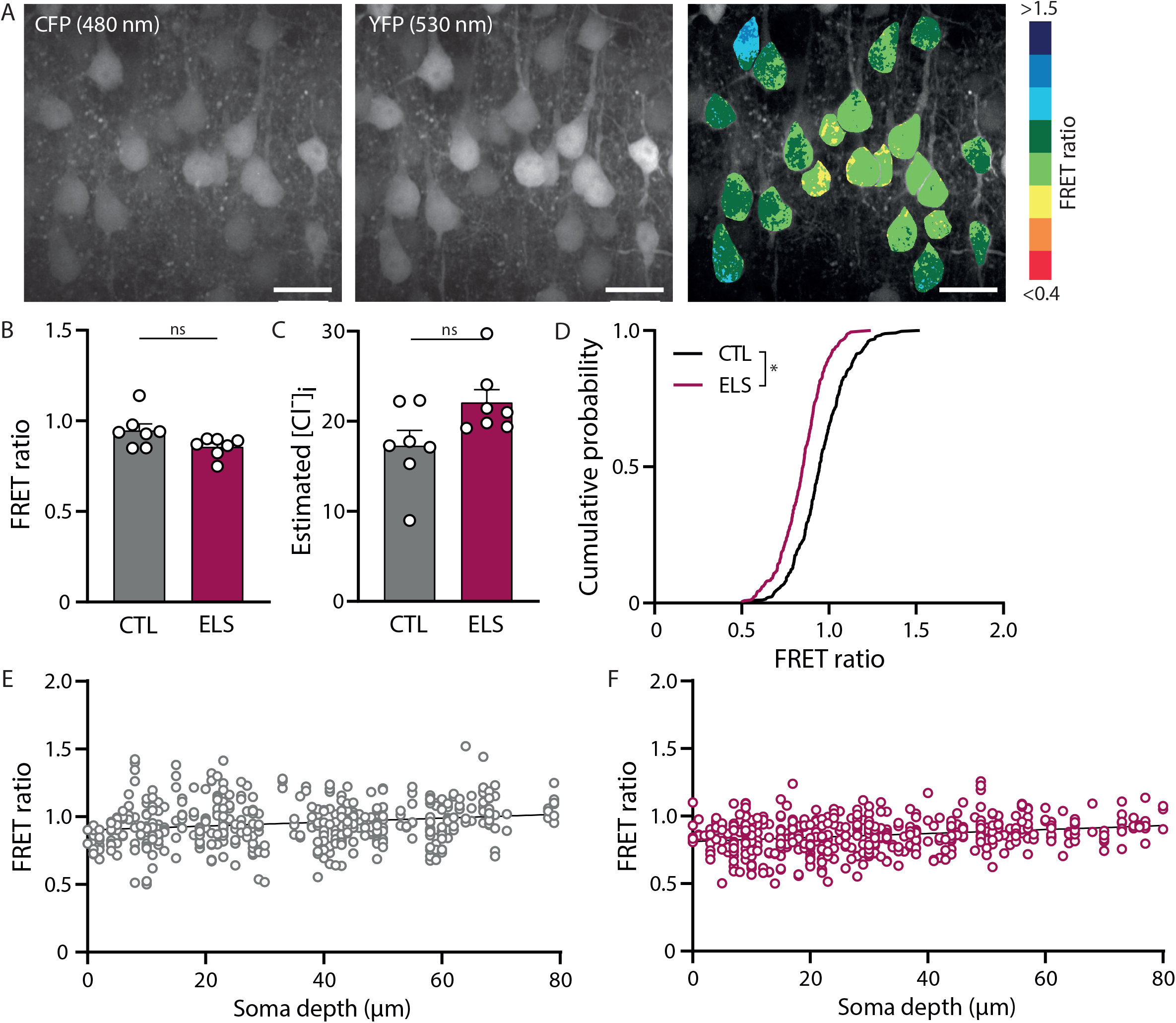
Higher [Cl^-^]_i_ in L2/3 cells of the mPFC from mice that experienced early life stress. A. Two-photon image of layer 2/3 neurons of the medial PFC of an acute slice of a P9 SClm mouse. Shown is the CFP and YFP fluorescence, and the corresponding FRET ratios. Individual cells are color-coded to their FRET ratios. Scale bar: 20 μm. B. Average FRET ratios for slice from control mice and from mice after ELS. Data from 7 mice in both groups (p = 0.55, T test). C. Average estimated [Cl^-^]_i_ as calculated from the FRET ratios in B (p = 0.07, MW test). D. Cumulative distribution of individual FRET ratios in slices from control and ELS mice. For each mouse 50 cells were randomly selected, making a total of 350 plotted cells per condition (p < 0.001; KS test). E. Individual FRET ratios plotted against soma depth in slices from control mice (n = 443 neurons). Line represents linear regression fit (r = 0.035; p < 0.0001). F. Same as E for slices from mice after ELS (n = 470 neurons). Line represents linear regression fit (r = 0.058; p < 0.0001).

### SClm sensor calibration

Calibrations for chloride were performed as described before (Boffi et al., 2018; Grimley et al., 2013; Rahmati et al., 2021). As SClm mice were no longer available we used organotypic hippocampal cultures from WT mice expressing the SClm sensor. Cultured slices were treated with ionophores (100 μM nigericin and 50 μM tributyltin acetate, Merck) to clamp [Cl^-^]_i_ and intracellular pH to extracellular levels. Saline containing various [Cl^-^]_i_ were perfused at approximately 1 mL/min. High chloride solution consisted of (in mM): 105 KCl, 48 NaCl, 10 HEPES, 20 D-glucose, 2 Na-EGTA, and 4 MgCl_2_, whereas the solution without chloride contained (in mM): 105 K-gluconate, 48 Na-gluconate, 10 HEPES, 20 D-glucose, 2 Na-EGTA and 4 Mg(gluconate)_2_. The high extracellular K^+^ concentrations are necessary for proper functioning of nigericin (Pressman and Fahim, 1982). Intermediate [Cl^-^]_i_ solutions (0, 5, 10, 50 and 100 mM) were prepared by mixing the two solutions. To maximally quench the SClm sensor we used a KF solution containing (in mM): 105 KF, 48 NaF, 10 HEPES, 20 D-glucose, 2 Na-EGTA and 4 Mg(gluconate)_2_. All calibration solutions were adjusted to pH 7.4. The first calibration solution with ionophores was washed in for 20 minutes and subsequent calibration solutions with ionophores were washed in for 15 minutes. Image z-stacks were acquired every 3 minutes at a resolution of 4.1 pixels/μm (512×512 pixels, 126×126 μm) with 1 μm steps of 30-50 μm in depth. FRET ratios reached a plateau after 10 minutes of wash in, which we assumed reflected equal intracellular and extracellular chloride concentrations. We constructed the calibration curve by plotting measured FRET ratios against extracellular chloride concentrations.

However, we noticed that FRET ratios differed widely between experiments, especially at intermediate chloride levels (5-10 mM). As this severely impaired robustness of the calibration in the most relevant chloride range, we resorted to perforated patch clamp measurements to calibrate the SClm sensor within the physiological range. We performed perforated patch clamp recordings in organotypic hippocampal cultures (described below) to determine [Cl^-^]_i_ in CA1 pyramidal cells at DIV1-3, 8-10 and 20-22. We plotted these values against the average FRET ratios measured in these cells at the same DIV and added these data to the calibration data from the ionophores. We fitted this composite calibration curve (Fig. 1D) with the following relation between FRET ratios and [Cl^-^]_i_ (Boffi et al., 2018; Grimley et al., 2013; Rahmati et al., 2021):

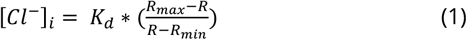

(in which *R* is the YFP/CFP emission ratio), to obtain the dissociation constant *K_d_* and the minimum and maximum FRET ratio *R_min_* and *R_max_* for our measurements.

### Electrophysiology

Whole-cell patch clamp recordings were made of pyramidal neurons in the hippocampal CA1 or mPFC of acute slices from SClm mice. Recording pipettes (resistance of 4-6 MΩ) were pulled from thick-walled borosilicate glass capillaries (World Precision Instruments) and filled with high chloride internal solution (in mM: 70 K-gluconate, 70 KCl, 0.5 EGTA, 10 HEPES, 4 MgATP, 0.4 NaGTP, 4 Na_2_-Phosphocreatine with pH 7.3 and osmolarity 295 mOsm/L). Cells were kept at a holding potential of −60 mV in voltage clamp throughout the experiment.

Perforated patch clamp recordings were made from CA1 pyramidal neurons in cultured hippocampal slices from WT mice at 30-32 °C. Recording pipettes (resistance of 2-4 MΩ) were pulled from thick-walled borosilicate glass capillaries (World Precision Instruments). The pipette tip was filled with gramicidin-free KCl solution (140 mM KCl and 10 mM HEPES, pH 7.2, and osmolarity 285 mOsm/L) and then backfilled with the KCl solution containing gramicidin (60 μg/ml, Sigma). CA1 neurons were clamped at −65 mV and the access resistance of the perforated cells was monitored constantly before and during recordings. An access resistance of 50 MΩ was considered acceptable to start recording. GABAergic currents were evoked by puffs of 50 μM muscimol (Tocris) dissolved in HEPES-buffered ACSF (in mM: 135 NaCl, 3 KCl, 2.5 CaCl_2_, 1.3 MgCl_2_, 1.25 Na_2_H_2_PO_4_, 20 Glucose, and 10 HEPES) in the presence of 1 μM TTX (Abcam). To determine the reversal potential of chloride, GABAergic currents were recorded at a holding potentials between −100 mV and −30 mV in 10 mV steps, upon local somatic application of the GABA_A_ receptor agonist muscimol (50 μM) dissolved in HEPES-buffered ACSF every 30s using a Picospritzer II. The GABA reversal potential was determined from the intersection of the current-voltage curve with the x-axis. We assumed that the chloride reversal potential *E_Cl_* equals the GABA reversal potential and used the Nernst equation to determine neuronal chloride concentrations:

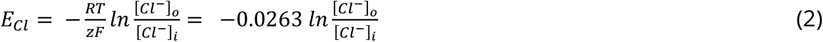

(with [*Cl*^-^]_*o*_ = 136.6 mM in ACSF) to convert the measured reversal potentials to estimated [Cl^-^]_i_. We are aware that GABA_A_ channels are also permeable for HCO_3_^-^ ions (Bormann et al., 1987; Kaila, 1994; Kaila et al., 1993). By assuming that *E_Cl_* equals the GABA reversal potential, we will slightly overestimate [Cl^-^]_i_.

### Statistical analysis

Statistical analysis was performed with Prism 9 (GraphPad). Normality was tested using Shapiro-Wilk tests. For unpaired samples statistical significance was evaluated using the unpaired Student’s t test (T test) for normally distributed data points, or the non-parametric Mann-Whitney (MW) test otherwise. A one-way ANOVA (or the Kruskal-Wallis (KW) test for non-normal distributions) was used when more than two groups were compared. Cumulative distributions were tested with the Kolmogorov-Smirnov (KS) test. Simple linear regression analysis was used to test if FRET ratios were influenced by the depth of the soma in the slice. P < 0.05 was considered significant. All data is presented as mean ± SEM.

## Results

### Two-photon imaging of [Cl^-^]_i_

To quantify the neuronal chloride concentration, SClm expression was targeted to pyramidal neurons by crossing SuperClomeleon^lox/-^ mice (Rahmati et al., 2021) with transgenic mice in which Cre recombinase expression was driven by the calcium/calmodulin-dependent protein kinase II alpha (CamKIIα) promoter (Casanova et al., 2001; Tsien et al., 1996). CaMKIIα is mostly expressed by excitatory neurons (Sík et al., 1998) and by some glia cells (Pylayeva-Gupta, 2011). In our slices from young mice, we observed many pyramidal neurons in the hippocampus and prefrontal cortex expressing the SClm sensor (Fig 1A). It is expected that ~35% of the neurons in the adult cortex and ~70% of the neurons in the adult hippocampus express CamKIIα (Wang et al., 2013). In our slices, the fraction of SClm cells is expected to be slightly lower, because CamKIIα expression still increases between P3 and P15 (Casanova et al., 2001).

The optogenetic SClm sensor consists of two fluorescent proteins, Cerulean (CFP mutant) and Topaz (YFP mutant), joined by a flexible linker (Fig. 1B). Binding of chloride to YFP reduces the FRET from the donor CFP to the YFP acceptor (Arosio and Ratto, 2014; Grimley et al., 2013). We used two-photon fluorescence microscopy to measure the 530 nm/480 nm (YFP/CFP) emission ratio (hereafter: FRET ratio) of individual cells. We calibrated measured FRET ratios against different [Cl^-^]_i_ using a combination of perforated patch recordings and ionophore treatment (Fig. 1C,D; see methods for details). Fitting this curve with equation (1) yielded a K_d_ value of 24.6 mM (Fig. 1D), which is in good agreement with previous reports (Grimley et al., 2013; Rahmati et al., 2021). The range of values for R_max_ measured in different calibration experiments was between 1.30 and 1.82. R_min_ ranged between 0.38 and 0.48. We used this fit to convert measured FRET ratios into [Cl^-^]_i_, for all our experiments, but we are aware that these should be considered reasonable estimates of the actual intracellular chloride levels at best.

### Acute manipulation of [Cl^-^]_i_ in brain slices

To assess the responsiveness of the SClm sensor to changes in intracellular chloride, we patched a pyramidal neuron in an acute hippocampal slice with a high concentration of chloride in the patch pipette, while monitoring changes in SClm FRET ratios. The FRET ratio in the patched cell changed immediately after break-in due to the rapid influx of chloride (Fig 2A,B). This decrease in FRET ratio was not observed in neighboring cells in the same field of view (Fig 2C). In a separate set of experiments, we monitored changes in FRET ratios during wash-in of muscimol, a specific GABA_A_ receptor agonist. As GABA_A_ receptors are chloride channels, activation of GABA_A_ receptors will induce influx of chloride ions, and therefore result in an increase in [Cl^-^]_i_. We observed a rapid 20% decrease in FRET ratio (indicating a 5-10 mM change in [Cl^-^]_i_) upon administration of muscimol, caused by the cellular influx of chloride (Fig 2D,E). These experiments demonstrate that the SClm sensor reliably reports rapid changes in neuronal [Cl^-^]_i_ within the physiological range.

### Development of neuronal [Cl^-^]_i_ levels in organotypic cultures

Next, we imaged neuronal [Cl^-^]_i_ levels in cultured hippocampal slices at different developmental stages. FRET ratios were determined at DIV2-3, DIV8-10 and DIV20-21 (Fig 3A). We observed a clear increase in FRET ratios, corresponding with a decrease in [Cl^-^]_i_, between DIV2-3 and DIV8-10 (Fig 3B,C). Although the average **FRET ratio was very similar** at DIV20-21 and DIV 8-10, the fraction of cells with high FRET ratios appeared larger at DIV20-21(Fig 3D),suggesting that chloride levels in some neurons were still **decreasing** at this age. FRET ratios were not dependent on the depth of the somata in the slice (Fig 3E-G). This indicates that cell-to-cell differences were not due to their location in the slice and confirmed that FRET values are independent of fluorescence intensity. Our results show that the GABA shift continues in organotypic hippocampal cultures during the first two weeks *in vitro.*

### Early life stress elevates neuronal [Cl^-^]_i_ at P9

We then used the SClm sensor to detect changes in chloride development in the prefrontal cortex in young mice after ELS. To induce ELS in young mice, we provided a limited amount of nesting and bedding material to the mothers between P2 and P9, resulting in fragmented and unpredictable maternal care (Karst et al., 2020; Naninck et al., 2015; Rice et al., 2008). To examine if this early life experience affected chloride maturation in the young pups, we measured FRET ratios in layer 2/3 neurons in acute slices from the mPFC from control and ELS male mice at P9, immediately after the stress period (Fig 4A). In total, 443 and 470 neurons were included in the analysis obtained from 7 control and 7 ELS mice, respectively. As expected, average [Cl^-^]_i_ levels were higher in mPFC compared to hippocampal pyramidal cells at comparable age (~17.3 mM in P9 mPFC slices (Fig. 4C) to ~11.5 mM in DIV2-3 cultured hippocampal slices (Fig. 3C)). This reflects the delayed maturation of the PFC compared to the hippocampus (Amadeo et al., 2018; Karst et al., 2019). The average FRET ratios in layer 2/3 pyramidal cells in slices from SClm mice that experienced ELS were similar compared to the average FRET ratios from control SClm mice (Fig 4B,C). However, when we analyzed the distribution of individual FRET ratios, we observed a significant shift towards more cells with a lower FRET ratio in the ELS condition (Fig 4D). No soma depth-dependent changes in FRET ratio were found in both conditions, indicating that FRET ratios could be reliably measured at variable depth in the slice (Fig 4E,F). Together our results show that ELS leads to an increase in neurons with high, immature chloride levels at P9 compared to control mice.

## Discussion

In this study we performed two-photon chloride imaging using the SClm sensor to determine the time course of chloride maturation in cultured hippocampal slices and to examine alterations of the chloride development in the mPFC by ELS. Previous non-ratiometric chemical indicators, including 6-methoxy-N-(3- sulfopropyl)quinolinium (SPQ) and N-(ethoxycarbonylmethyl)-6methoxyquinolinium bromide (MQAE) (Illsley and Verkman, 1987; Verkman et al., 1989), have the disadvantage that their fluorescence depends not only on [Cl^-^]_i_, but also on the dye concentration and optical thickness at each location, e.g. depth in the slice or in the brain. Therefore SPQ and MQAE allow for the assessment of acute changes in [Cl^-^]_i_ within the same neurons (Arosio and Ratto, 2014; Zajac et al., 2020), but cannot be used to study developmental changes which requires comparisons between animals and between slices. **The** SClm sensor has an improved affinity for chloride compared to its processor Clomeleon (Berglund et al., 2006), resulting in a more than fourfold improvement in signal to noise over Clomeleon (Grimley et al., 2013). Two-photon chloride imaging poses major advantages over perforated patch clamp recordings. Most importantly, chloride imaging allows for assessing of [Cl^-^]_i_ over time in multiple neurons simultaneously in a non-invasive manner. One important limitation of the SClm sensor is its sensitivity to intracellular pH (pH_i_) (Lodovichi et al., 2022). However, we do not expect large changes in pHi in our in vitro experiments, and pH_i_ remains fairly constant during postnatal development (Sulis Sato et al., 2017). Using SClm, we could directly measure changes in [Cl^-^]_i_ when we loaded 70 mM chloride into a neuron via the patch pipette and after addition of a GABA_A_ receptor agonist. This demonstrates that the SClm sensor reliably reports changes in intracellular chloride within the physiological range. Furthermore, we could detect subtle changes in the distribution of individual [Cl^-^]_i_ levels within the pyramidal cell population during normal and disturbed postnatal development. Subtle changes at the population level are physiologically relevant and would have been hard to pick up otherwise.

Although the advantages of direct chloride imaging using the SClm sensor are numerous, we found that the conversion of FRET ratios to absolute values of intracellular chloride concentrations was not very robust. In our hands, calibration using ionophores and varying extracellular chloride concentrations gave variable results. Unreliable FRET ratios, especially in the physiological range (5-10 mM), hampered reliable calibration to [Cl^-^]_i_. Compared to our cultured slices, calibration in cultured neurons are better accessible by ionophores and changes in extracellular chloride levels due to the 2D structure (Boffi et al., 2018; Grimley et al., 2013). In addition, strong regulation of intracellular chloride levels (Kaila et al., 2014; Rahmati et al., 2021) and variable resilience of neurons to the harsh calibration conditions may have hampered the calibration procedure in our slices. We therefore resorted to using perforated patch clamp and inferred chloride concentrations from the reversal potential of GABA_A_ currents. Although our calibration curve was in good agreement with previous reports (Grimley et al., 2013; Rahmati et al., 2021), we noticed that small alterations in the fit strongly affect chloride level estimates. We therefore conclude that the SClm sensor is an excellent tool to measure relative changes in [Cl^-^]_i_ which are physiological relevant, but that conversion to absolute chloride concentrations should be interpreted with care. In our view, this disadvantage does not outweigh the significant benefits of using the SClm sensor to detect changes in neuronal chloride levels over time and between conditions. In recent years, genetically encoded fluorescent sensors have been developed which enable imaging of different molecules. Live imaging studies using these sensors, most prominently of intracellular calcium, have made great contributions to our understanding of intra-and intercellular signaling (Day-Cooney et al., 2022; Dong et al., 2022), despite the fact that calibration of most of these sensors to absolute concentrations remains notoriously difficult. We hope that current and future chloride sensors (Lodovichi et al., 2022; Zajac et al., 2020) will make a similar impact on our understanding of chloride homeostasis.

Using the SClm sensor, we have monitored the developmental decrease in neuronal chloride levels in organotypic hippocampal cultures from SClm mice. The SClm sensor proved much more sensitive than its precursor Clomeleon, which was previously used to determine chloride maturation in cultured neurons and in an in vitro epilepsy model (Dzhala and Staley, 2021; Kuner and Augustine, 2000). With the SClm sensor we could discern that neuronal chloride levels in our cultured slices show a clear reduction between DIV3 and DIV9 (equivalent to the second postnatal week *in vivo)*, and that in some pyramidal cells chloride levels continue to decrease until DIV22. The large cell-to-cell differences that we observe have been previously reported (Dzhala et al., 2012; Kirmse et al., 2015; Stein et al., 2004; Sulis Sato et al., 2017; Yamada et al., 2004). The estimated [Cl^-^]_i_ values in our developing cultured slices are in good agreement with previous estimates in cultured hippocampal neurons (Kuner and Augustine, 2000; Tyzio et al., 2007), and acute hippocampal slices (Staley and Proctor, 1999; Tyzio et al., 2007). A recent study using LSSmClopHensor chloride sensor in the visual cortex *in vivo* (Sulis Sato et al., 2017) also showed a rapid reduction in [Cl^-^]_I_ during the first postnatal week, followed by a slow further decrease to mature levels.

Brain development is strongly influenced by external factors and early life experiences (Hensch, 2004; Miguel et al., 2019). Here we used ELS, an established model to interfere with early brain development with long-lasting consequences for psychopathological risks later in life (Bachiller et al., 2022; Catale et al., 2022; Joёls et al., 2018; Teicher et al., 2016). We used SClm to detect possible alterations in chloride maturation in the mPFC. We observed that ELS results in a shift towards higher (i.e. immature) chloride levels in individual layer 2/3 cells in the mPFC. This suggests that ELS delays the GABA shift in SClm mice, but we did not examine chloride levels at older ages. Our results are in line with previous reports showing a delayed GABA shift in hippocampal neurons after prenatal maternal restraint stress and when newborn pups were repeatedly separated from their mother (Furukawa et al., 2017; Hu et al., 2017; Veerawatananan et al., 2016). However, the effect on the GABA shift can be very sensitive to the type and timing of stress, and a different maternal separation paradigm results in an advance of the hippocampal GABA shift (Galanopoulou, 2008). Using the same ELS paradigm in C57/BL6 mice, we previously reported accelerated maturation of synaptic currents in layer 2/3 mPFC pyramidal cells (Karst et al., 2020), and decreased [Cl^-^]_i_ levels in young pups after ELS (Karst et al., 2019). Although we used the same stress paradigm and mice were housed in the same facility, the effect of ELS on [Cl^-^]_i_ was remarkably different between the SClm and WT mice. Chloride homeostasis is highly regulated and affected by many intracellular factors including the chloride buffering capacity (Rahmati et al., 2021), and we cannot exclude that the permanent presence of a chloride sensor induces subtle changes in the regulation of chloride homeostasis in neurons in SClm mice. However, the [Cl^-^]_i_ we report here in SClm (unstressed) controls (17.3 mM) was comparable to the chloride concentration that was previously measured by perforated patch in WT C57/BL6 control mice (18.8 mM) (Henk Karst, unpublished observations), suggesting that the difference cannot be explained by a difference in baseline chloride levels. The response to stress can differ substantially between mouse strains and C57BL/6 mice appear more resilient to stress in comparison to other strains (Murthy and Gould, 2018). Unfortunately, we did not directly compare the two mouse lines. A difference in ELS response may indicate a difference in genetic predisposition with possible consequences for stress responses and stress-related behavior (Joëls et al., 2018; McIlwrick et al., 2016; Teicher et al., 2016). The poor breeding performance of SClm mice may also reflect differences in stress response compared to C57BL/6 mice. Our results underscore that the developmental chloride trajectory appears incredibly sensitive to environmental factors and may differ between mouse lines. We advise to take these differences into account in behavioral experiments.

Our data demonstrate that the high sensitivity of the SClm sensor at relevant chloride concentrations allows detecting physiological alterations in neuronal chloride levels during normal and altered postnatal development. Although we also found has some limitations, our study underscores that two-photon chloride imaging is a powerful technique to further illuminate the role of chloride signaling in the brain.

## Acknowledgements

We thank Prof. Kevin Staley for providing the SuperClomeleon^lox/-^ mouse line and SCIm AAV and Dr. Stefan Berger for the CamKIIα^Cre/-^ mice. We thank several members of the Staley lab for helpful discussions. We thank René van Dorland for his technical support with the AAV. We thank Prof. Marian Joёls for her constructive feedback on the manuscript.

